# Quantitative Learning of Cellular Features From Single-cell Transcriptomics Data Facilitates Effective Drug Repurposing

**DOI:** 10.1101/2023.09.16.558051

**Authors:** Jianmei Zhong, Junyao Yang, Yinghui Song, Zhihua Zhang, Chunming Wang, Renyang Tong, Chenglong Li, Nanhui Yu, Lianhong Zou, Sulai Liu, Jun Pu, Wei Lin

## Abstract

In this study, we have devised a computational framework SuperFeat that allows for the training of a machine learning model and evaluate the canonical cellular states/features in pathological tissues that underlie the progression of disease. This framework also enables the identification of potential drugs that target the presumed detrimental cellular features. This framework was constructed on the basis of an artificial neural network with the gene expression profiles serving as input nodes. The training data comprised single-cell RNA-seq datasets that encompassed the specific cell lineage during the developmental progression of cell features. A few models of the canonical cancer-involved cellular states/features were tested by such framework. Finally, we have illustrated the drug repurposing pipeline, utilizing the training parameters derived from the adverse cellular states/features, which has yielded successful validation results both *in vitro* and *in vivo*. SuperFeat is accessible at https://github.com/weilin-genomics/rSuperFeat.

## Introduction

Since the emergence of various high-throughput -omics technologies such as microarray [1] and next-generation-sequencing (NGS) [2], researchers have been generating vast amounts of molecular profiling datasets of the biological and clinical samples at an unprecedented rate. The single-cell sequencing technique has even added another degree of magnitude and a new dimension of information [3–6]. Many models and algorithms have been developed to evaluate the biological signal and characterize the sample. Nonetheless, most of the existing methods are hardly generalizable but specific to particular problems [7,8]. A universal framework for the rapid yet generic assessment of the biological/clinical samples using molecular profiling data could be very useful and efficient, especially for the -omics assays at high granularity.

Human learning involves the processes of classification and quantification. The artificial neural network (ANN) has been used as a flexible framework to simulate and streamline such processes using a machine/computer. It provides a relatively simple but generalizable classification and quantification model to automatically structure the human knowledge acquired from a large body of data. The ANN nodes of the neural network reflect the qualitative or the quantitative state of a biological concept in the human mind. Most importantly, subsequent human decisions, such as clinical decisions, could be made based on the evaluation of the cellular state. Therefore, the ANN structure could be used to realize the automatic learning and evaluation of biological state/feature and thus facilitate making decisions more efficiently based on the input of a high volume of data from the high-throughput -omics techniques. This mechanism is becoming increasingly powerful and can accomplish many unprecedented tasks, even in the biomedical field.

Previously, we developed the ANN-based cell-type classifier framework, SuperCT [9]. It uses the single-cell RNA-seq digital expression profiles as input to characterize the canonical cell lineages. Similar cell-type classification strategies have been published since then [10,11]. For the first time, the ANN-based classifier was used in the machine learning of high-dimension single-cell expression data to solve the cell-type classification problem. In a fully connected neural network for a cell-type classifier, the node weight reflects the empirical contribution of a gene to a certain cell type which also make this classifier interpretable.

In the meantime, a certain cell type could undergo a spectrum of variable cellular states reflected by a few function and signaling-related gene expressions. For instance, T cells turn into an exhaustion state in most solid tumors [12]; macrophages undergo polarization in certain biological contexts [13,14]. Nonetheless, assessing the variability of such cellular state by a couple of markers using the current barcoded-bead (BCB) based single-cell transcriptomics data oftentimes was not robust. This is due to the stochastic nature of the RNA transcription within a single cell [15] and the limitation of detection for a specific molecule by the current barcoded-bead-capture [4]. Unless we perform a more comprehensive assessment using a group of coordinately expressed genes, the state of a single cell might not be accurately evaluated. The good news is that a certain cellular state/feature is always associated with a group of up- and down-regulated genes, thus can be reflected by a scoring strategy based on the presence/absence of a group of transcriptions.

The strategy of activating or reversing a certain cellular state of the disease-involved cell type could be employed for possible therapeutics. For example, immune checkpoint inhibition has been used in cancer therapy. It serves the purpose of reversing the T cell exhaustion [16]. Targeting helper T cellular state conversion could be used in mitigating autoimmunity [17]. Idally, if small molecules could be used to reverse the adverse cellular state or promote the beneficial cell state. Such an idea was implemented in the Connectivity Map (CMap) bulk (but not single-cell) transcriptomics datasets. Cell lines have been used to generate the CMap perturbagen transcriptomics data repository, which was used to search for repurposing drugs [18,19]. Nonetheless, prior to the application of the scRNA-seq technique to clinical samples, the CMap search was oftentimes compromised by the convoluted signals of a mixture of cell types of different roles. The high-resolution transcriptomics data like single-cell RNA-seq allows to characterize the target cell population and thus makes the CMap search more specific. SignatureSearch provided a flexible tool to use ranked gene set to search perturbagen database that could be used to achieve such goal [20].

In addition to the single-cell RNA-seq, a similar BCB-based RNA quantification strategy is applied in the 10× Genomics Visium Spatial transcriptomics platform. Obviously, the cellular state/feature evaluation strategy could be applicable to this type of data and provide a very user-friendly spatial visualization of the histological slides. The landscape of the cellular state/feature will be smoothed even with the stochastic transcriptions of the functional genes on the spots.

In this study, we aimed to build an ANN-based framework in order to efficiently perform the learning and quantitative assessment of a variety of biological features documented in the literature. This framework can facilitate the thorough evaluation of the critical features of the tissue samples or even the precise diagnoses of the patients with adequate details. Gene-set-based methods such as Seurat modular score [21], AUCell [22], singscore [23] and GSVA [24] can somewhat do a similar job. We plan to extensively compare their performance in order to understand the strength of the new method.

## Results

### An efficient framework of feature learning and scoring using single-cell transcriptomics

We constructed a fully connected ANN, called Supervised Feature Learning and Scoring (SuperFeat), to learn the quantitative features with one-single node in the hidden layer and two output nodes. The value of the middle-layer node will be used for the quantitative evaluation of the cellular state/feature. **Figure 1** illustrates the workflow of the entire framework. Figure 1A shows a single-cell dataset contains multiple cell types. If one of the cell types (circled by dashed line) was annotated with a variable cellular state/feature (according to some signature genes) and can be manually divided into two annotated subpopulations, this variable cell population could be used to train the SuperFeat model for evaluating this cellular state/feature. If there are more datasets with the similar cellular state annotation, we should include the cell populations from more datasets to enhance the representativity and the model generalizability. To enable the users to customize their own model, we also provided the python code (Figure 1B) for those who likes to train their own cellular state/feature model and to assess similar transcriptomics data. The Figure 1B also shows the structure of the ANN and the green node basically provides score to evaluate how much the transcriptional profile should be categorized to state 1 (in red) or state 0 (in black).

**Figure 1.**
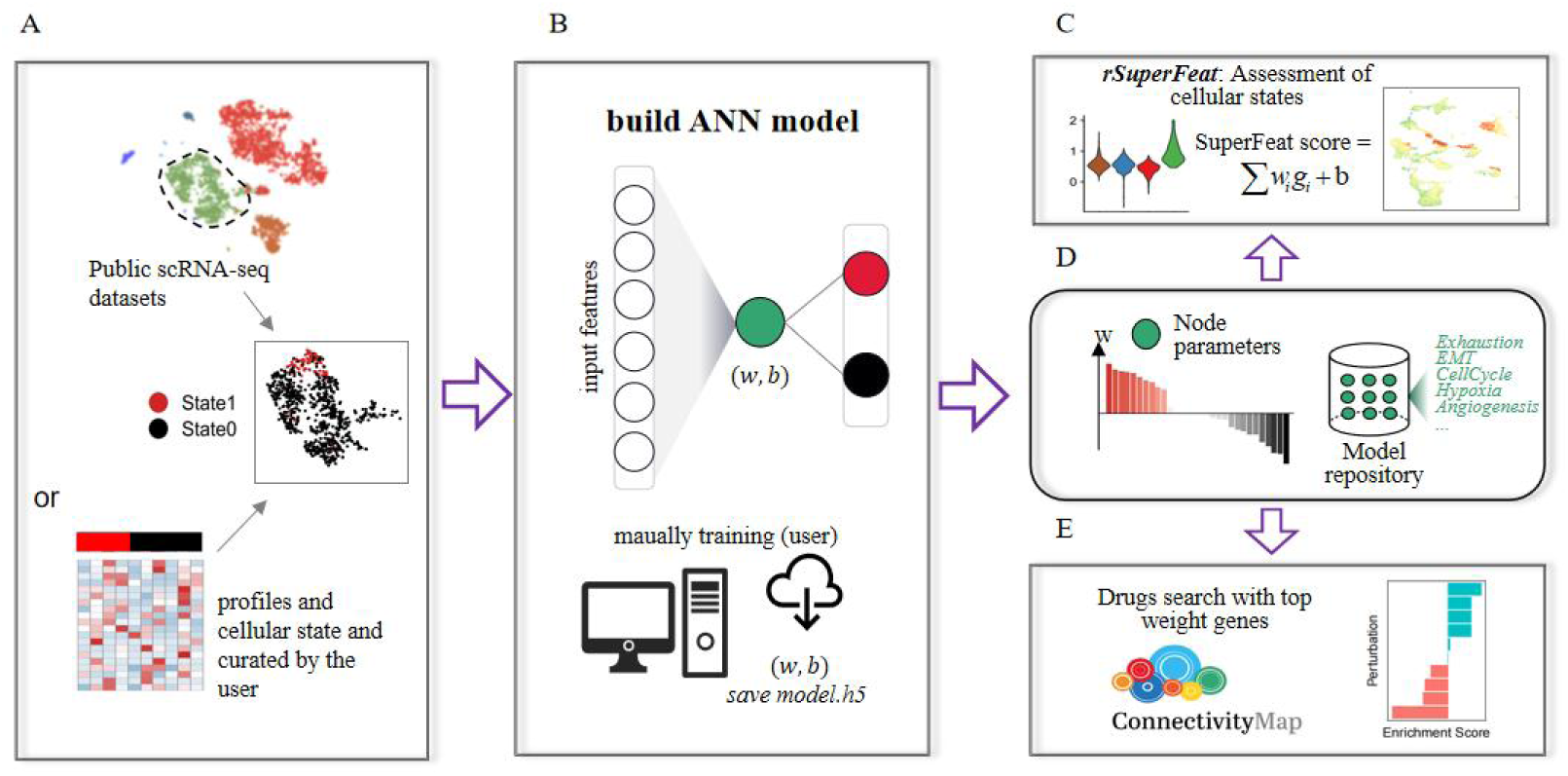
Workflow of SuperFeat framework. **A**. A diagram showing how the cell lineage with variable cellular state/feature is defined in the training dataset. **B**. ANN structure and implementation. **C**. The parameters of the models trained from dataset and the repository of models for the canonical cellular states/features. **D**. Evaluation of the cellular states/features using SuperFeat model and visualization. **E**. Drug search using the model parameters. ANN, artificial neural network.

Figure 1C shows the weights of the genes contribute to the score in this ANN model. It also shows how a repository was organized for the models trained from the literatures for a group of canonical cellular states and features [https://github.com/weilin-genomics/SuperFeatModels]. Users are also welcome to contribute their own models to this repository if they have a convincing cellular state hypothesis and their paper is published. Figure 1D shows how the model could be used to calculate the score of the cellular state/feature of the cells from another dataset of single-cell RNA-seq or spatial RNA-seq in the applicable biological context. This part of application was implemented in an R package ‘rSuperFeat’ (https://github.com/weilin-genomics/rSuperFeat). Later, as shown in Figure 1E, the model parameters can be further utilized to search for a Connectivity Map (CMap) [18] perturbagen that potentially enhances or alleviates the stress associated with cellular state change.

### Training and evaluation of four tumor hallmark cellular states/features

Using our most recent version of SuperFeat training framework and the training dataset from a study of kidney renal clear cell carcinoma [25], we were able to first perform the training of the T cell exhaustion model parameters and apply the scoring model to assess the T cells. The exhaustion scores allowed us to discern the exhausted and active T/NK cell populations that was correlated to the immunotherapy. The distributions of the exhaustion scores of the annotated populations of T cells in training dataset were shown as **Figure 2A**. The layout of these cells is shown in **Figure S1**. The canonical marker signals of exhaustion annotated by the authors are shown as **Figure 2B**, confirming the overall concordance. Then we used the datasets of another study including the infiltrating T cells from hepatocellular carcinoma samples published recently [26,27] to test the model performance. The *C4*-*CD8*-*LAYN* population stands out (**Figure 2C**), which is concordant to the paper’s cluster interpretation. The layout of the cell populations in the testing datasets and T cell exhaustion scores are shown in **Figure S2A-B**.

**Figure 2.**
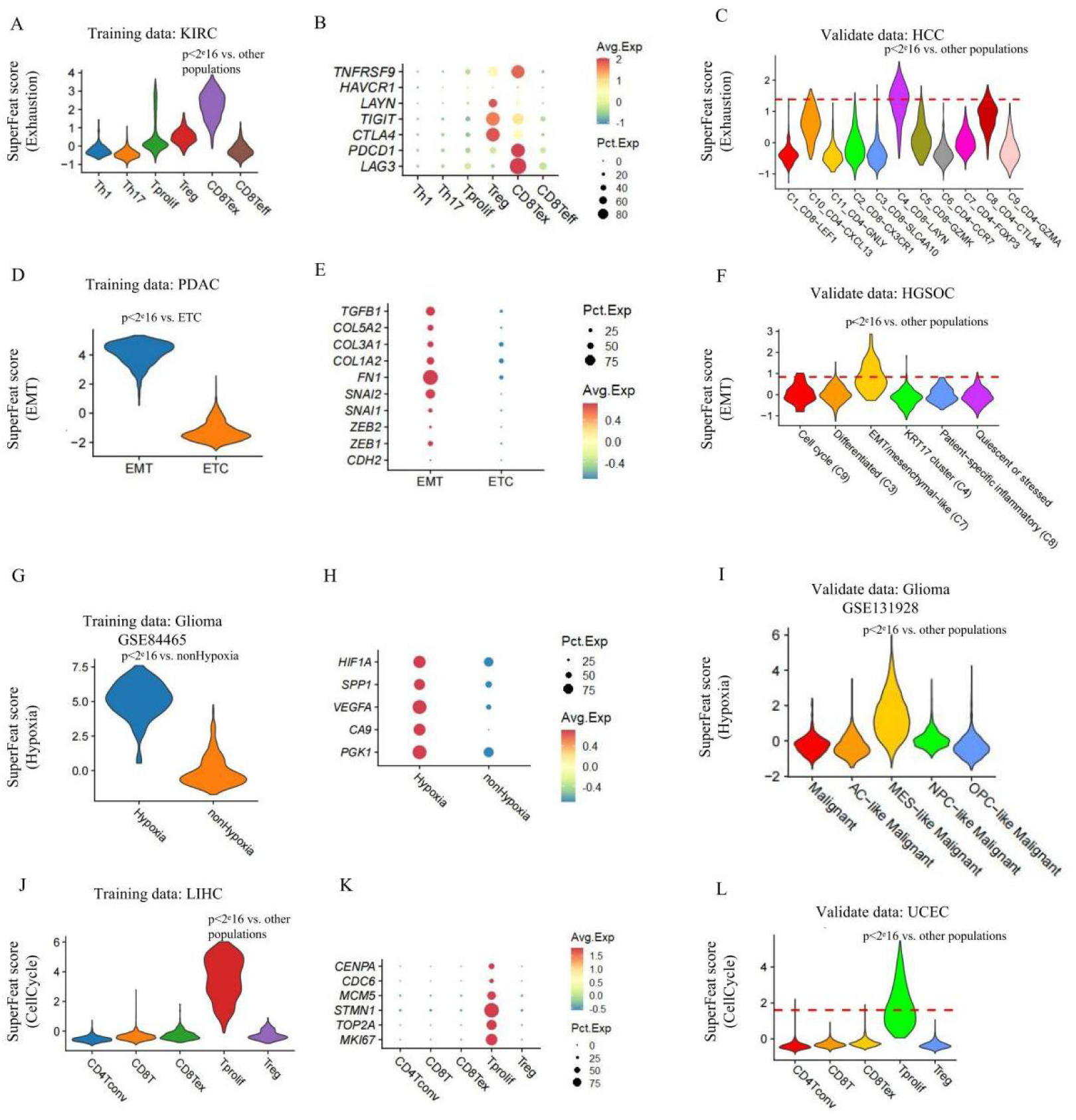
The tumor related cellular states/features defined by SuperFeat. **A.** T cell exhaustion SuperFeat scores on training data. **B**. T cell exhaustion markers. **C**. SuperFeat scores validated by a HCC dataset. **D.** EMT SuperFeat scores on training data. **E**. Several canonical EMT markers. **F**. SuperFeat scores validated by a HGSOC dataset. **G.** Hypoxia SuperFeat scores on training data. **H**. Hypoxia markers. **I**. SuperFeat scores validated by a Glioma dataset. **J.** CellCycle SuperFeat scores on training data. **K**. CellCycle markers. **L**. SuperFeat scores validated by a UCEC dataset. All significance was determined by Wilcoxon test. HCC, hepatocellular carcinoma; HGSOC, high-grade serous ovarian cancer; UCEC, uterine corpus endometrial carcinoma.

Epithelial mesenchymal transition (EMT) is another example of the variable cellular state that oftentimes occurs in the epithelial cells under wound healing, organ fibrosis, and, especially initiation of the metastasis of cancer progression. In here, we used a set of pancreatic ductal adenocarcinoma single cell data with EMT annotation in the tumors to train a EMT model [28]. We used the EMT population from high-grade serous ovarian tumor to test the performance of this model [29]. The scores of the training cells are shown as **Figure 2D** and the scores of the testing cells are shown as **Figure 2F**. The C7 clusters, which was annotated as ‘EMT/mesenchymal-like’ population exhibit higher EMT signals. Both show concordance with the published annotation and the canonical EMT markers (**Figure 2E**).

Hypoxia refers to a state in when the cells are deprived of adequate oxygen. Tumors oftentimes develop such a hypoxic feature. We used Darmanis *et.al.* Dataset [30] to train the hypoxia model and used Neftel *et.al.* Dataset [31] to evaluate the feature. The results are shown in **Figure 2G-I**. The annotated hypoxic cell populations (MES-like malignant clusters) in the testing dataset gave high SuperFeat scores and exhibit canonical hypoxic gene expressions such as *HIF1A*, *VEGFA* etc.

The proliferation rate is essential to the development of tissue. For example, the immunohistochemistry (IHC) signature Ki67 (with gene symbol *MKI67*) has been commonly used to evaluate cell proliferation by pathologists. Here, we also trained a model of proliferation [32] and tested this model performance [33]. **Figure 2J and L** show the proliferation scores in both training and testing data and Figure 2K shows the transcriptional signals of the canonical CellCycle genes.

### SuperFeat model parameters uncover involved known and novel features

The neural network used to be considered a black-box learning machine. As our framework has a very simple network structure with a limited number of nodes in the middle layer and the input genes were trained independently, the contribution of the input nodes (genes) can be easily evaluated by the weight values. Therefore, the top positive and the top negative weight genes could be examined and performed enrichment analysis in order to understand how they underlie the cellular state change.

For T cell exhaustion model trained from Neal *et.al.* Dataset [25], the canonical marker genes such as *PDCD1*, *LAG3* and *HAVCR2* etc. shows high-rank weight with no surprise (**Figure 3A**). The enrichment analysis based on the top positive weight genes of T cell exhaustion hits ‘regulation of cell activation’ (**Figure 3B**). The enrichment analysis based on the top negative genes of T cell exhaustion shows lymphocyte activation. Obviously, they are two polarized states of T cells.

**Figure 3.**
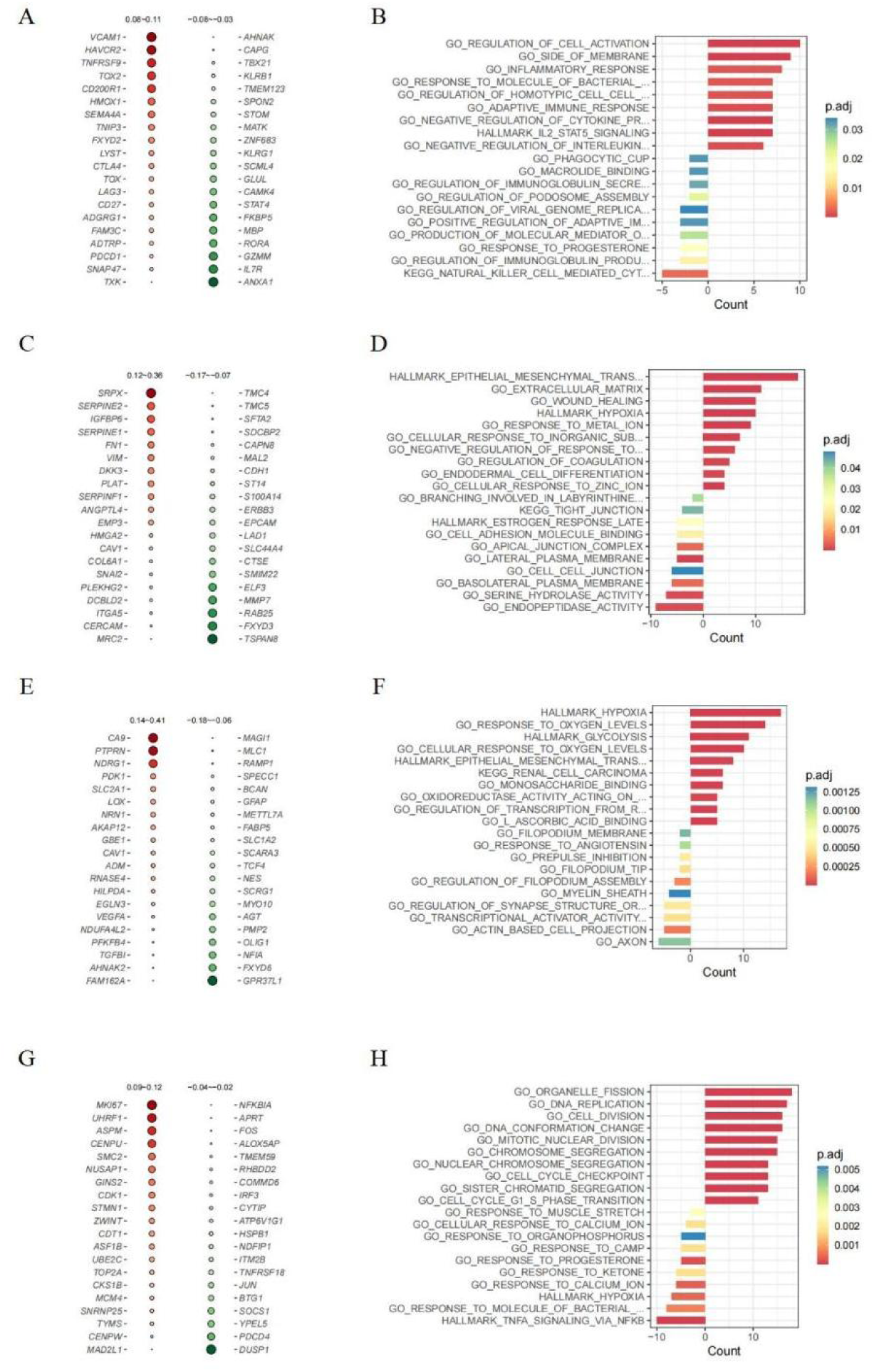
Interpretation of SuperFeat Model parameters. **A**. Top 20 weighted genes of T cell exhaustion model. **B**. Enrichment analysis on the top 50 positive/negative weighted genes of T cell exhaustion model. **C**. Top 20 weighted genes of EMT model. **D**. Enrichment analysis on the top 50 positive/negative weighted genes of EMT model. **E**. Top 20 weighted genes of hypoxia model. **F**. Enrichment analysis on the top 50 positive/negative weighted genes of hypoxia model. **G**. Top 20 weighted genes of CellCycle model. **H**. Enrichment analysis on the top 50 positive/negative weighted genes of CellCycle model. Negative counts indicate enrichments for negative weighted genes and the positive counts indicate positive ones.

For EMT model trained by Lin *et.al*. Dataset [28] the top positive weight genes include *SERPINE1*, *VIM*, *SNAI2,* and *ANGPTL4*, which are also expected (**Figure 3C**). The top positive genes exactly hit the hallmark of EMT (**Figure 3D**). The top negative weight genes of EMT hits endopeptidase activity, which has never been discussed and probably needs extra attention.

For hypoxia model trained from Darmanis *et.al*. Dataset [30], *VEGFA*, *CAV1* and *CA9* etc. emerged at the top positive weight genes (**Figure 3E**), which are canonical hypoxia-related genes. The top positive genes exactly hit the term of ‘hallmark of hypoxia’, which again validates the reliability of our framework (**Figure 3F)**. Interestingly, the enrichment analysis based on the top positive weight genes of hypoxia also hits the hallmark of glycolysis. This correlation has been reported validated in the previous study [34]. Such correlation couldn’t be found using gene-set-based scoring systems.

For CellCycle model trained from Zhang *et.al.* Dataset [32], *MKI67*, *STMN1*, *MCM4* and *TOP2A* etc. showed up at the top positive weight genes (**Figure 3G**), which are canonical cell cycle-related genes. These top positive genes hit the terms such as ‘Cell Division’, ‘Cell cycle checkpoint’ etc. (**Figure 3H**).

In summary, the top positive weighted genes are mostly involved in the pathways that drive the development of the corresponding cellular states/features, and the top negative genes are involved in signature genes of the opposite cellular states. It makes the ANN-based model parameters interpretable.

### Outperformance of SuperFeat over gene-set-based scoring in discerning the target subpopulations

Prior to SuperFeat, people oftentimes used Seurat modular scoring, AUCell, singscore and GSVA etc. to evaluate the cellular state based on a group of canonical genes. To efficiently discern the cell subpopulations of a certain cell type that reflect the variability of a certain cellular state/feature, the separation of the signal distributions of the scores suggests a better performance. In here, we demonstrated the heatmaps of median signal scores (**Figure 4A-D**). It appeared that SuperFeat scores make the target subpopulation stand out with better color contrast than others. We further used the Kolmogorov-Smirnov statistics to numerically evaluate how much the subpopulations with certain cellular states/features stand out from their counterpart subpopulations without this feature. The results show that SuperFeat mostly outperforms other scoring systems by the relatively larger Kolmogorov-Smirnov D value (**Figure 4E,G,I,K**) and larger area-under-curve (AUCs, shown as **Figure 4F,H,J,L**), except being slightly lower than the Seurat scores for EMT gene-set. More detailed results of the accuracy analysis are shown in **Table S1**.

**Figure 4.**
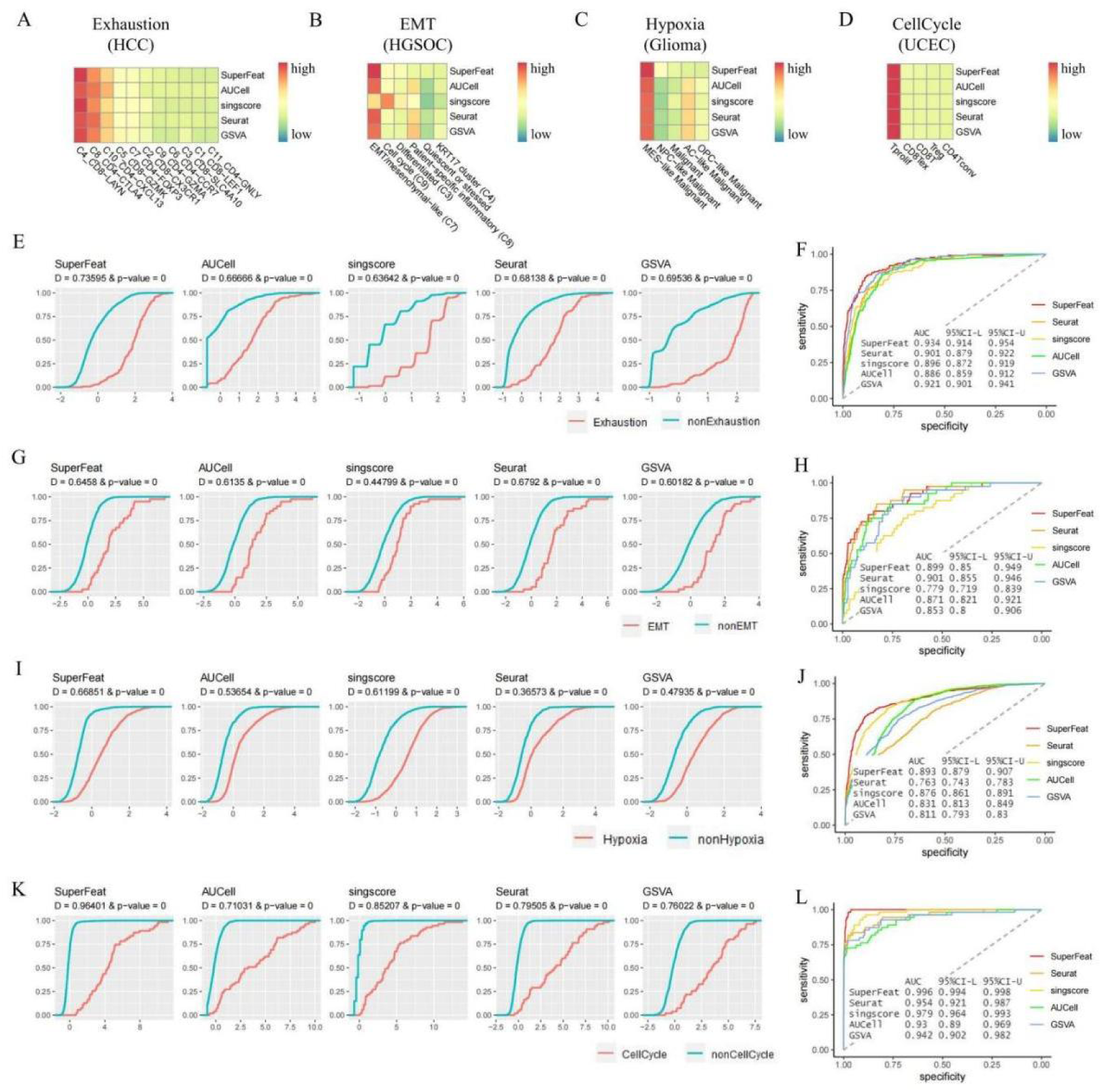
Comparison of cellular state/feature scoring methods. **A**. Heatmap for the SuperFeat scores of T cell exhaustion. **B**. Heatmap for the SuperFeat scores of EMT. **C**. Heatmap for the SuperFeat scores of hypoxia. **D**. Heatmap for the SuperFeat scores of CellCycle. **E**. K-S statistics on T cell exhaustion state. **F**. AUC analysis on T cell exhaustion state. **G**. K-S statistics on EMT state. **H.** AUC analysis on EMT state. **I.** K-S statistics on hypoxia state. **J**. AUC analysis on hypoxia state. **K**. K-S statistics on CellCycle state. **L**. AUC analysis on CellCycle state.

Overall, it is confident to claim, SuperFeat discerns the clusters annotated with the designated cellular states/features more easily and accurately than other gene-set-based scoring methods. Furthermore, the advantage of SuperFeat scoring resides in its independency of the canonical gene set, which saves the arbitrariness of gene selection.

### Feature scoring in spatial transcriptomics study

The SuperFeat scoring models can also be applied to the spatial transcriptome data, which allows us to correlate the cellular states/features to the histology. We used the two 10× genomics Visium slides from the tumor samples of a cholangiocarcinoma cancer patient to demonstrate the mapping of the proliferative signals. **Figure 5A** shows the original H&E staining of the two slides from the same tumor tissue. **Figure 5B** shows the area of the pathological annotations of the two slides determined by an experienced pathologist. **Figure 5C** shows the landscape of the proliferative signal on the two slides. **Figure 5D** shows the violin plots of the proliferative SuperFeat scores of the two slides. From the two replicate slides of the same tissue, the high cell cycle signal is reproducibly enriched in the immune regions, suggesting the high proliferative potential. We also mapped the signal of the two signature genes, *MKI67* and *TOP2A* on the 10× genomics Visium slides (shown as **Figure 5E-F**). The landscape of the two genes’ signals appears more stochastic than SuperFeat proliferation signal, suggesting the advantage of a comprehensive evaluation. The other three tumor-related feature scores are shown in **Figure S3**.

**Figure 5.**
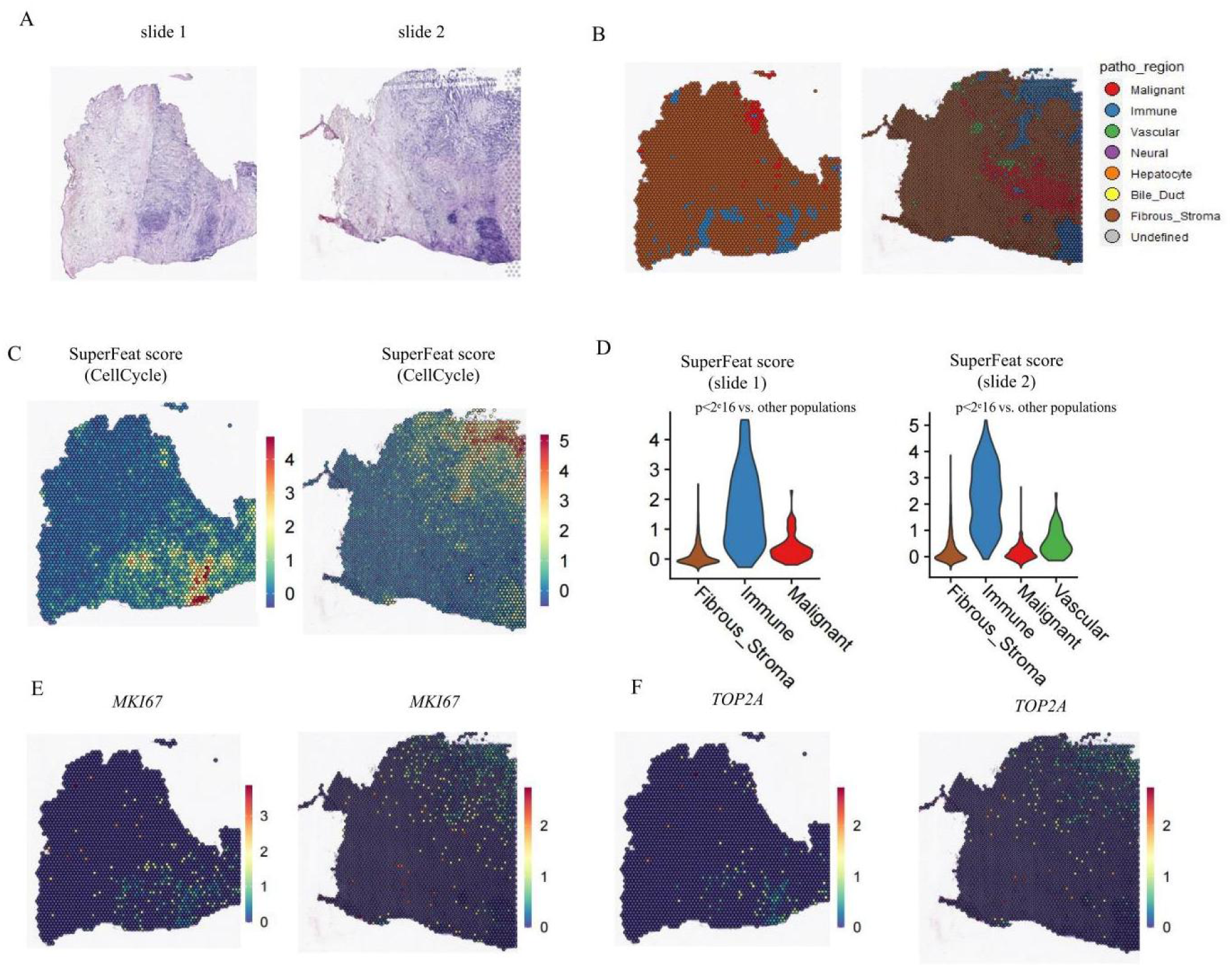
SuperFeat scoring on two spatial transcriptomics slides. **A**. H&E staining for two slides from the same tumor tissue. **B**. histological annotation by the pathologist. **C**. SuperFeat CellCycle scores of the dots on the two replicate slides of 10×genomics Visium platform. **D**. Violin plots of SuperFeat CellCycle scores of the dots on the two replicate slides. The significance was determined by Wilcoxon test. **E**. *MKI67* signals of the dots on the two replicate slides. **F**. *TOP2A* signals of the dots on the two replicate slides.

### Repurposing drug search using SuperFeat model parameters

As the malignant cellular states such as tumor proliferation and EMT could be targeted for cancer therapy, similar to CMap strategy, we were able to perform the search based on the gene weights in the trained cellular state model against the perturbagen databases such as CMap and LINCS L1000. Without confounding the variability derived from the heterogeneous composition of the cell populations in the samples, we intuitively believed that the drug search using the top weight-ranked genes derived from single-cell data could be more specific to the cellular state change than the differential genes found in bulk RNA study but more general in the application targeting the detrimental cellular state.

Using CMap search strategy over the LINCS L1000 data repositories implemented by the signatureSearch package of R [20], we tested such an idea and got interesting results.

**Figure 6A** shows the drug search based on the top-weighted genes derived from the cell cycle feature. Among the top 10 hits on LINCS, NVP-BEZ235 [35], AZD8055 [36], palbociclib [37], naproxol [38], ivermectin [39], oxindole-I [40], are the CellCycle-arrest agents that has been previously reported.

**Figure 6.**
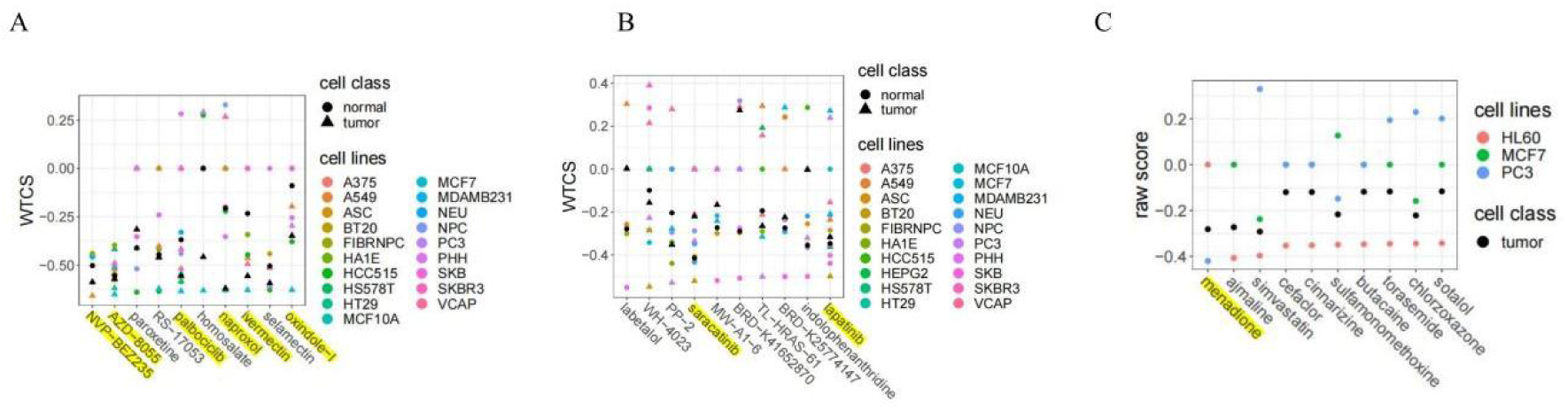
Drug search using weight-ranked genes of SuperFeat. **A**. Drug search result based on the top positive/negative weighted genes of the SuperFeat CellCycle model and LINCS L1000 database. **B**. Drug search results based on the SuperFeat EMT model and LINCS L1000 database. **C**. Drug search results based on the SuperFeat EMT model and CMap database.

The drug search based on the top-weighted genes derived from the EMT feature also gives meaningful results. Among the top 10 hits on LINCS, saracatinib has been reported with the restoration of E-cadherin expression [41] and the mesenchymal-epithelial-transition role of lapatinib was reported [42] (**Figure 6B**). The top 1 hit on CMap, menadione (Vitamin K3) has been reported with suppression of EMT [43] (**Figure 6C**).

### Stability of model parameters and reproducibility of SuperFeat results

As the model parameters are the major determinants of the drug search, in order to evaluate the robustness of the methods, we compared the top weighted genes and the output drugs derived from the different training datasets using the most established cellular feature, *i.e.*, proliferation.

The datasets were retrieved from Gene Expression Omnibus database: GSE140228 (hepatocellular carcinoma proliferation) and GSE110686 (T cell proliferation), respectively. Although different cell types were involved in the same cellular state, our results show 55-60% of the top positively weighted genes are reproducible in two datasets (statistics shown in the Venn diagram of **Figure 7A**). There were also 21-28% overlapped negatively weighted genes, whose roles remain elusive. The enrichment analysis of the top positively weighted genes mostly hit the same GO terms (**Figure 7B**). At last, we compared the output drugs. For CMap search, 7 out of 20 top selections are reproducible. For LINCS L1000 search, 10 out of 20 top selections are reproducible (**Figure 7C**). It was also found that naproxol, palbociclib, NVP-BEZ235 were always among the top selections of both outputs.

**Figure 7.**
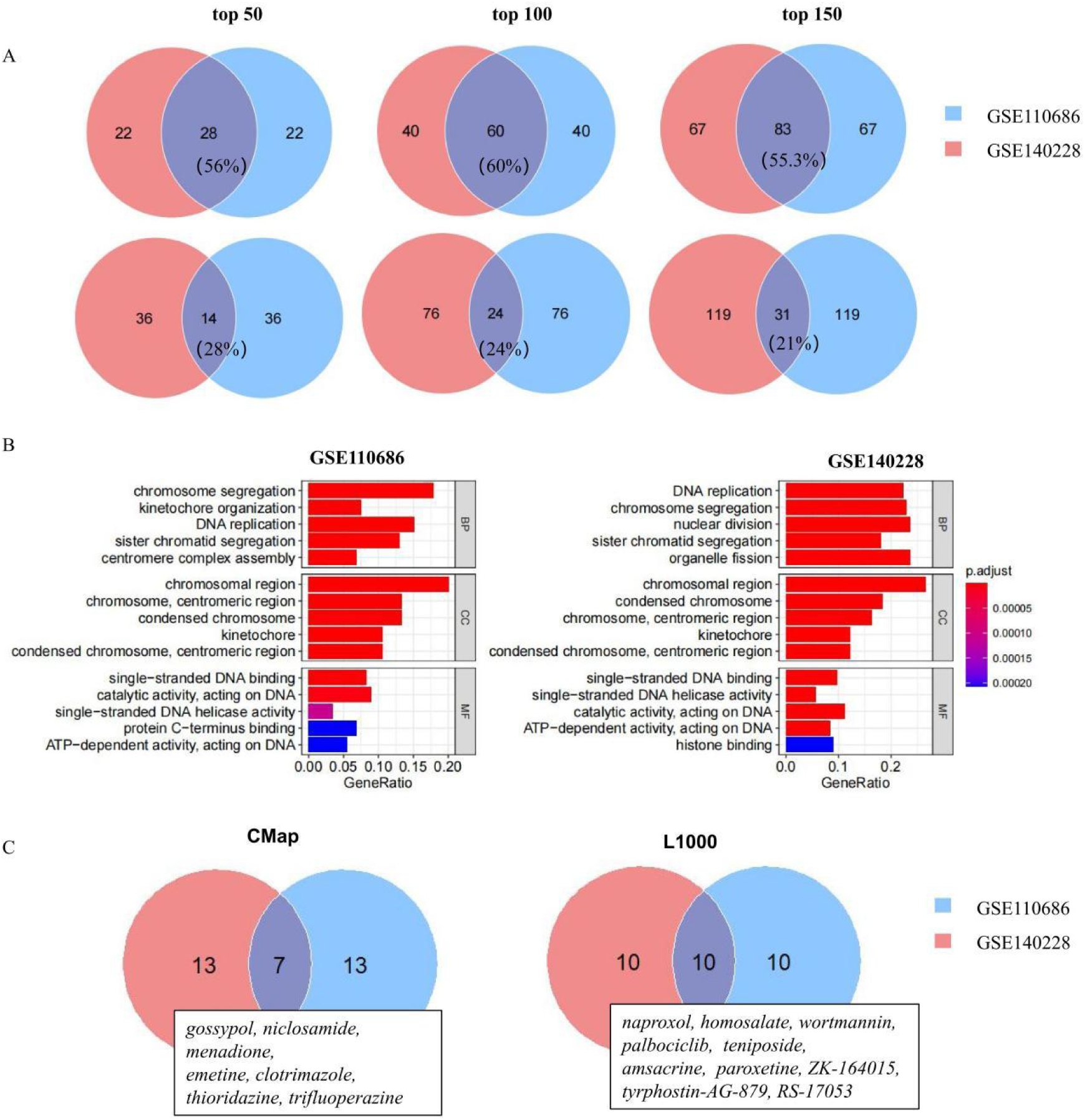
Stability of model parameters and reproducibility of SuperFeat results. **A**. Venn diagrams show stable weighted genes between different datasets. top panel: positively weighted genes, bottom panel: negatively weighted genes. The number indicates genes count. **B**. The barplot display similar GO terms. only top 5 categories are shown here. **C**. Venn diagrams show drugs was repeatedly searched and detailed drugs are listed. the number indicates drugs count.

Such reproducible results suggest that, despite the training datasets being different, similar genes that contribute to the canonical cellular state/feature will be assigned high weights. These genes will also subsequently provide the leads necessary to discover drug candidates that either promote beneficial cellular states or inhibit adverse ones.

### *In vitro* and *in vivo* validation of drug effect on adverse cellular state development in tumor

In a collaborative study of the cancer-associated-fibroblast (CAF), we discovered a state transition of CAF in a subcutaneous mouse tumor model. We have proved a Cd34^+^Pi16^+^ fibroblast progenitor gives rise to the classical Acta2^+^ Mcam^+^ CAF in tumor, also known as myCAF [paper under submission, personal communication]. Similar CAF subpopulations have been characterized in both human and mouse tissues in the published datasets [44] (**Figure 8A-B**). The developmental trajectory of CAF from the Cd34^+^ fibroblast progenitor state to the Acta2^+^ classical CAF state was defined and validated in our collaborative study. State #1 (Cd34^+^ CAF) represents a more stemness state characterized by the signature genes such as *Cd34* and *Pdgfra*. State #0 (Cd34^-^ CAF) represents a more differentiation state with the tumor-prone signature marker such as α-SMA (**Figure 8C**).

**Figure 8.**
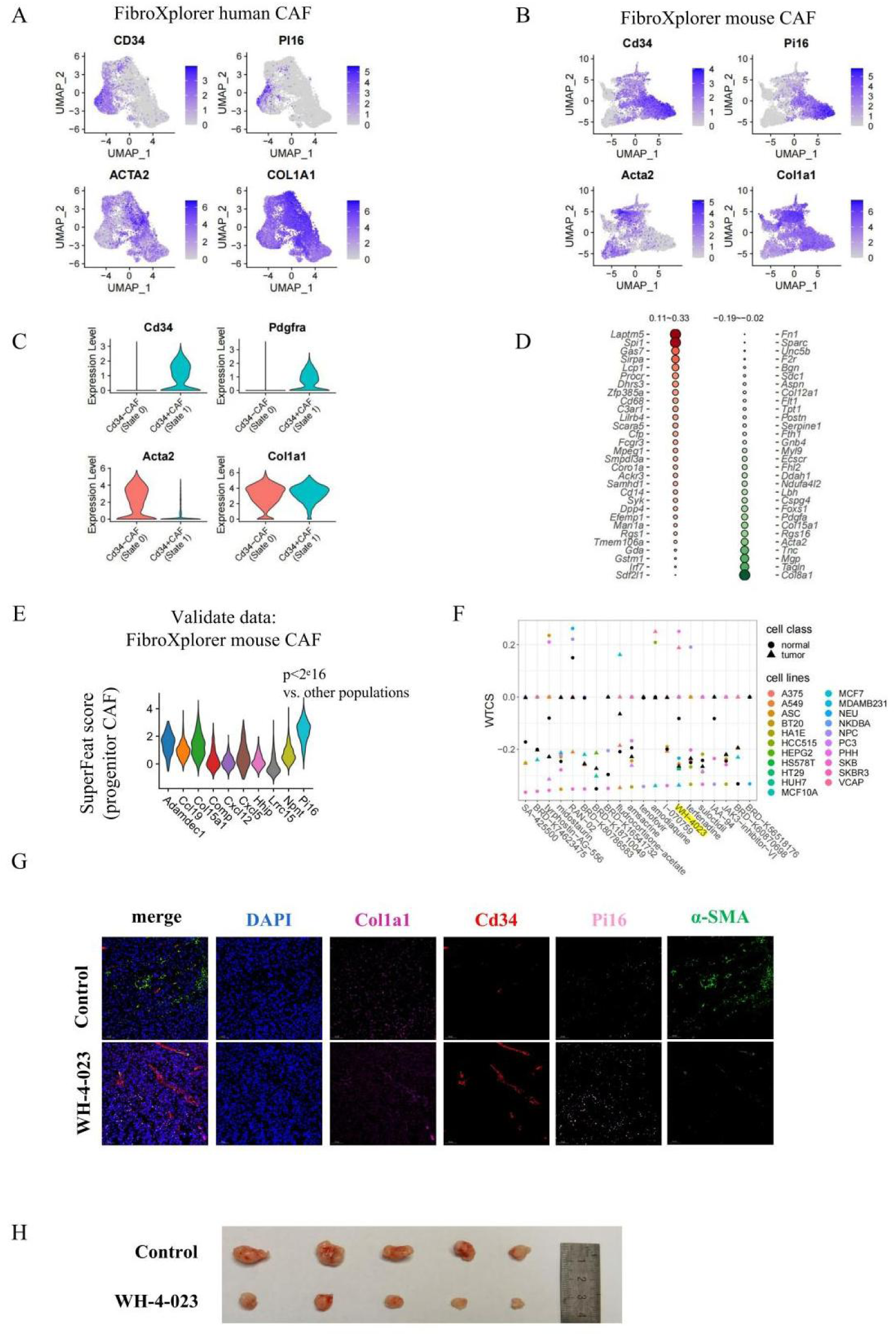
*in vitro* and *in vivo* validation of a drug candidate by SuperFeat and SignatureSearch. **A**. Fibroblast subpopulations in human tissues from FibroXplorer database. **B**. Fibroblast subpopulations in mouse tissues from FibroXplorer database. **C**. Violin plot for the fibroblast subpopulations in our CAF development study in subcutaneous tumor mice. **D**. SuperFeat model top 50 parameters for CAF development. **E**. Validation of CAF development SuperFeat model using FibroXplorer mouse dataset. **F**. Top 20 drug search candidates. **G**. The effect of WH-4-023 on fibroblast development using IF. **H**. The effect of WH-4-023 on tumor shrinkage. The significance was determined by Wilcoxon test. FibroXplorer, https://www.fibroxplorer.com/home.

Using SuperFeat framework, we were hence able to train a new scoring model to calculate the progression of CAF differentiation in the tumoral biopsy. Using the subsequent search, we aimed to find the drugs that potentially suppress the conversion from the progenitor State #1 to the tumor-prone State #0 (**Figure 8D-E**). The top 20 drug candidates from LINCS L1000 are shown in **Figure 8F**. By validation experiments on these 20 candidates, four of them showed very promising outcomes (detailed results in another paper under submission). **Figure 8G** shows the outcome of MKN-45 (gastric cancer cell) subcutaneous tumor model with the treatment of WH-4023, which is one of the 4 drugs. The tumors significantly shrank in 5 replicates.

To confirm that the drug indeed inhibited tumor growth via the suppression of CAF development, we performed immunofluorescence (IF) staining for signature genes, including Cd34/Pi16 for the progenitor state, and α-SMA/Collagen for the classic tumor-prone state. We extracted CAFs from tumor tissue and cultured them in a differentiation medium (B16) mixed with tumor cell culture medium in a 3:1 ratio. We added the drug candidate directly to the differentiation medium, changing the medium every other day. After 14 days, we compared the IF signals and found that myCAF development was indeed curtailed. Notably, the WH-4023 treatment group exhibited lower α-SMA/Collagen signals but higher CD34/PI16 signals in comparison with the control group (**Figure 8H**)

## Conclusion

In this study, we’ve established and demonstrated an innovative framework designed to assess cellular states, tissue development, and even individual patient conditions. This framework comprises a few critical elements: A. an R package that facilitates the evaluation of canonical cellular states via a scoring system. B. a data repository containing model parameters of various documented cellular states from previous research. C. Python training code for user-defined cellular states. This setup allows for the efficient evaluation of canonical cellular states previously identified in research. Users can leverage this framework to simply assess cellular states using existing models, or train their own SuperFeat scoring models and potentially propagate their own cellular state models.

We’ve specifically demonstrated how the SuperFeat model assesses several key cellular states or features of tumors, such as cell proliferation, T cell exhaustion, hypoxia, and EMT - all of which are hallmark cancer signals. These examples underscore the simplicity, generalizability, and efficiency of the new method. We strived to demystify the ‘black box’ of the SuperFeat ANN model by visualizing and interpreting the model parameters, specifically the gene weights potentially contributing to a specific cellular state. Compared to the signature gene-set-based scoring method, SuperFeat demonstrated superior performance because it considers not only the up-regulation of signature genes, but also suppressed genes. Such contributions are reflected by positive and negative weights, respectively. By examining the weights of genes, researchers may acquire additional insights.

Essentially, SuperFeat provides another learning framework and a scoring strategy that streamline the rapid assessment of states or features in tissue, many of which have been extensively explored and acknowledged in existing literature. Moreover, compared with other traditional methods, SuperFeat renders such assessment more intuitive, generalizable, and user-friendly, without relying on the details of the gene set. While an Artificial Neural Network (ANN) model bypasses the uncertainties of statistical assumptions in a rigorous mathematical model, the weight parameters of genes derived from such a training still align remarkably well with expectations.

Logically, the metric values of the SuperFeat cellular state/feature reflect the resemblance between the putative cells-of-interest in the training dataset and the cells in the new study, in comparison to their counterparts. Therefore, the model’s reliability largely depends on the quality of the training dataset. As our understanding of cells and single-cell transcriptomics datasets keep growing, the SuperFeat framework is set to make the automated learning and assessment of cellular states/features increasingly reliable and efficient.

An even more thrilling advantage of this framework lies in its potential for downstream applications, *e.g.*, repurposing drug searches. The other gene-set-based approaches of cellular state evaluation are not comparable in this aspect. As we know, CMap was based on the connectivity between gene expression profiles and drug perturbations. Previous CMap perturbation datasets were derived from bulk RNA-seq . The recent progress of the single-cell RNA-seq technique makes it practical to generate such connectivity at single-cell resolutions. The advantage of our strategy could heighten the precision and success rate of expression-profile-based drug searches. We successfully conducted the validation experiments that substantiate the feasibility of such strategy. Another advantage of neural-net-based strategy pertains to the convenience of the interconnection with other models. It makes integration of drug search task based on multi-data sources possible.

In brief, our effort aims to build a preliminary and universal framework that allows to rapidly and automatically evaluate the conditions of multi-cellular samples by investigating the single-cell transcriptomics and eventually find the solutions for biological and clinical problems.

## Materials and methods

### Organization and preprocessing of the training and testing dataset

The datasets for feature training were summarized in **Table S2**. The cell populations with the designated cellular states/features came from the unsupervised clustering and enriched the canonical markers for the specific state/feature. For the training dataset, we only included the cell types in which the states/features were supposed to be exhibited in the model training. 19,202 genes (shown in **Table S3**) were taken into the input. The input genes were derived from total genes of the msigdbr v7.4.1 gene-set of subcategory H and C5 [45]. The missing expression values in the training or the testing dataset of these 19, 202 genes will be fill up with zero. The gene expression values of the digital expression matrix were rescaled to 0-1.

### Implementation of the SuperFeat artificial neural network

The artificial neural network structures and the learning strategy were implemented using Keras API. We designed a fully connected ANN model for the cell-type classification. Similar to what has been done for SuperCT, the inputs were transformed into the binary values of 19, 202 genes. As seen in most of the flow-cytometry analyses, the presence/absence of the signature gene partially contributes to the state/feature of the cell. Also, the binary signal input is compatible across most of the Unique-Molecular-Index-based (UMI) scRNA-seq platforms with the robust performance of the cell-type classification.

The input layer is connected to a hidden dense layer with only one hidden layer neuron using ReLU (Rectified Linear Unit) activation functions. The input layer has the L1 regularization with coefficient of 0.01. The output layer has two nodes. To avoid the under-representation of the small-sample-size cell types in the calculation of the accuracy function, we include the class-weight based on the sample size of each type in the model training. The loss function was defined as categorical cross-entropy.

### Evaluation of the designated features using SuperFeat Score

The cellular state/feature was evaluated using the following equation 1.

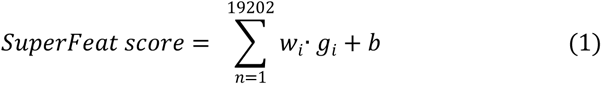

*w_i_* denotes the weight values from the model trained from the training dataset to the first layer. *g_i_* denotes whether the gene is detected in this cell or not. *b* denotes the bias value for the single node of the first hidden layer.

### Drug search using weight-ranked genes

We used qSig function of R package signatureSearch to perform the connectivity ranking based on the perturbagen × cell gene expression signatures over the CMap and LINCS L1000 databases. Top 250 positive and negative weighted genes were output by the printTopWeights function of rSuperFeat package and input to signatureSearch for the potential drugs. The rank of the drugs was based on the overall positive or negative connectivity scores, suggesting it could drive or reverse the change of the cellular state/feature, respectively.

### Unsupervised clustering, dimensional reduction, and data visualization in single-cell RNA-seq analyses

For the single-cell RNA-seq dataset downloaded from Gene Expression Omnibus database, the unsupervised clustering, dimensional reduction and data visualization in this paper were realized by a widely used scRNA-seq analytical suite, Seurat [46,47]. The Seurat objects were generated for each dataset with their digital expression matrices as input. The PCA was performed by Seurat RunPCA function. The tSNE coordinates were calculated using Seurat RunTSNE function. The UMAP coordinates were calculated using Seurat RunUMAP function. The putative clusters were defined by Seurat FindClusters function using the top 10 principal components and other default parameters. If the cell annotation were provided by the author in the database, we will directly use their annotation. If not, the unsupervised clusters were re-annotated according to the enriched literature markers.

### Cell Culture

The GC cell line MKN-45 cell was purchased from the Cell Bank of Type Culture Collection of the Chinese Academy of Sciences and cultured in Dulbecco’s Modified Eagle’s Medium (DMEM; Gibco, Life Technologies) with 10% FBS and antibiotics (100 IU/mL penicillin and 100 µg/mL streptomycin). The cell culture was placed in humidified air at 37°C with 5% CO2/95% air (v/v).

### *In vivo* subcutaneous tumor and drug treatment

In vivo studies, 4-6-week-old male BALB/c nude mice were purchased from Shanghai Laboratory Animal Center of China. Specifically, animals were housed in a controlled environment with a 12 h light/dark cycle, with free access to water and food and at temperatures of 21–23 °C and humidity of 40– 60%.

Low passage MKN-45 cells were resuspended in a 1:1 mixture of PBS and Matrigel (Corning #356 231) at 1 × 106 cells /mL. 100 µl of cell stock was injected subcutaneously on the shaved right flank of BALB/c nude mice. After10 days’growing, the tumor volume increased to an average size of 60 mm3. WH-4023 (Selleck, S7565) dissolved in DMSO(D2650, Sigma-Aldrich,). WH-4023(0.5mg/kg) was injected into the tumors every other day for 6 days. The same volume of DMSO was included as control. The mice were examined at regular times until they were sacrificed. The tumor size was measured using a digital caliper, and the tumor volume was calculated with the following formula: volume = 0.5 × width2 × length.

All animal breeding, housing and experimentation were conducted according to the guidelines of the Institutional Committee of Shanghai Jiao Tong University School of Medicine for Animal Research (Approval No. XHEC-F-2020-026).

### Immunofluorescence staining

The tumor tissues were harvested and fixed in 4% paraformaldehyde, embedded in paraffin, followed by cryosection with a thickness of 5–10 μm. The cryosections were fixed with 4% paraformaldehyde, permeabilized with 0.1% Triton X-100 in PBS for 15 minutes, and blocked with 10% FBS/PBS for 1 hour. Then stained with The primary antibodies: anti-CD34 (ab81289, Abcam), Pi16(PA5-111740, Invitrogen), Col1a1(Invitrogen, PA5-29569), a-SMA(ab124964, Abcam) antibody. The primary antibodies were diluted in 10% FBS/PBS by the dilution factor recommended by the suppliers, applied to the samples, and incubated at 37°C for 1.5 hours or at 4°C overnight. The secondary antibodies were diluted at 1:1000 in 10% FBS/PBS, then applied and incubated at 37°C for 45 minutes. The cell nucleus was counterstained with DAPI at room temperature for 5 minutes, and coverslips were mounted on slides with fluorescent mounting media.

## Code availability

The framework is implemented in R package rSuperFeat and can be accessed at https://github.com/weilin-genomics/rSuperFeat.

## CRediT author statement

**Wei Lin**: Conceptualization, Methodology, Supervision, Writing - original draft, Writing -review and editing. **Jianmei Zhong**: Methodology, Investigation, Visualization, Writing - review and editing. **Junyao Yang**: Investigation, Visualization, Writing - review and editing. **Yinghui Song**: Investigation, Visualization. **Zhihua Zhang**: Investigation, Visualization. **Chunming Wang, Renyang Tong, Chenglong Li, Chenglong Li, Lianhong Zou, Sulai Liu**: Investigation. **Jun Pu**: Supervision, Writing - review and editing.

## Acknowledgments

This study was supported by grants from Shanghai Jiaotong University, Renji Hospital Start-up funding for New PI, Young Talent of Hunan (2020RC3066), Hunan Natural Science Fund for Excellent Young Scholars (2021JJ20003), China Postdoctoral Science Foundation (2021T140197).

