## Supplementary material for "Quantitative Learning of Cellular Features From Single-cell Transcriptomics Data Facilitates Effective Drug Repurposing": Figure S1-3; Table S1-2

### Supplementary Figures

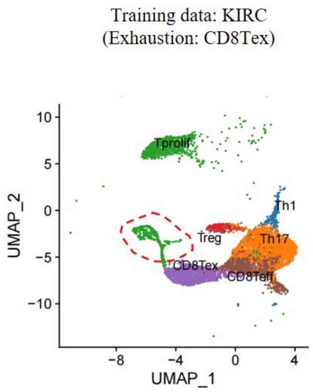


**Figure S1. UMAP layout of the training dataset for T cell exhaustion model**

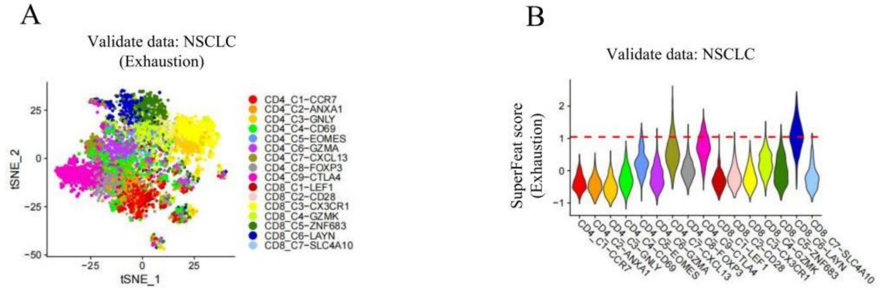


**Figure S2. T cell annotations in two published datasets.**

**A.** UMAP layout of T cell exhaustion in NSCLC validation dataset. **B**. Violin plot of T cell exhaustion SuperFeat scores. NSCLC, non-small-cell lung cancer.

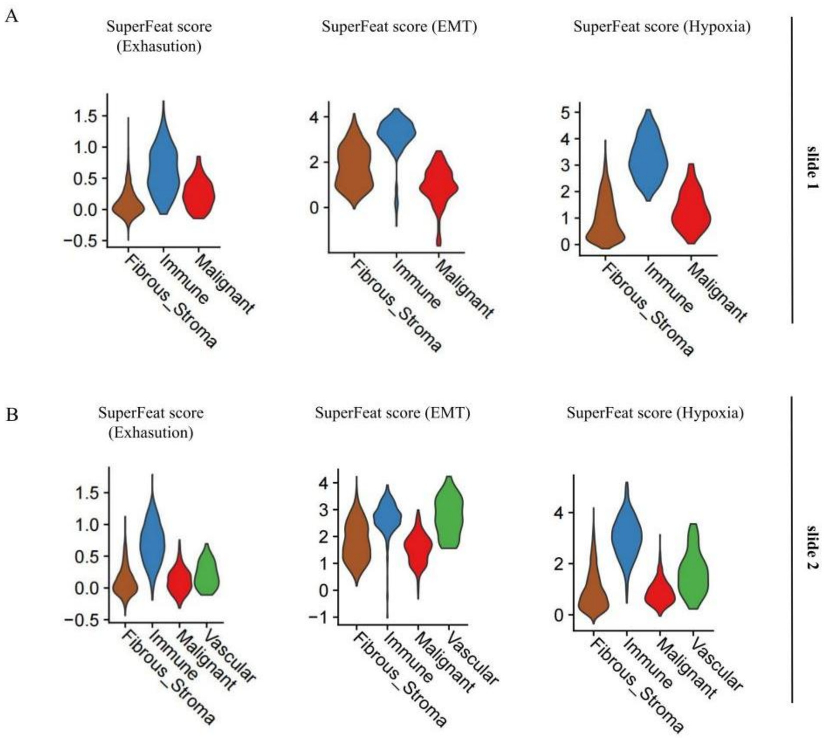


**Figure S3. SuperFeat feature scores based on the spatial transcriptomics.**
**A.** Violin plots for T cell exhaustion, EMT, and hypoxia signals on slide 1. **B**. Violin plots for T cell exhaustion, EMT, and hypoxia signals on slide 2.

### Supplementary Tables

**Table S1.** Outperformance of SuperFeat over gene-set-based scoring in discerning the target subpopulations

| ***Exhaustion*** | **AUC** | **95%CI-L** | **95%CI-U** | **threshold** | **specificity** | **sensitivity** | **accuracy** |
|---|---|---|---|---|---|---|---|
| SuperFeat | 0.934 | 0.914 | 0.954 | 0.782 | 0.885 | 0.851 | 0.883 |
| Seurat | 0.901 | 0.879 | 0.922 | 0.159 | 0.809 | 0.872 | 0.814 |
| singscore | 0.896 | 0.872 | 0.919 | 0.539 | 0.863 | 0.773 | 0.857 |
| AUCell | 0.886 | 0.859 | 0.912 | 0.141 | 0.808 | 0.858 | 0.812 |
| GSVA | 0.921 | 0.901 | 0.941 | 0.097 | 0.823 | 0.872 | 0.827 |
| ***EMT*** | | | | | | | |
| SuperFeat | 0.899 | 0.850 | 0.949 | 0.343 | 0.871 | 0.775 | 0.868 |
| Seurat | 0.901 | 0.855 | 0.946 | 0.113 | 0.829 | 0.850 | 0.830 |
| singscore | 0.779 | 0.719 | 0.839 | 0.239 | 0.673 | 0.775 | 0.676 |
| AUCell | 0.871 | 0.821 | 0.921 | 0.078 | 0.764 | 0.850 | 0.766 |
| GSVA | 0.853 | 0.800 | 0.906 | 0.030 | 0.752 | 0.850 | 0.755 |
| ***Hypoxia*** | | | | | | | |

| | | | | | | | |
|---|---|---|---|---|---|---|---|
| SuperFeat | 0.893 | 0.879 | 0.907 | 0.450 | 0.884 | 0.784 | 0.831 |
| Seurat | 0.763 | 0.743 | 0.783 | 0.032 | 0.561 | 0.805 | 0.692 |
| singscore | 0.876 | 0.861 | 0.891 | 0.235 | 0.786 | 0.826 | 0.808 |
| AUCell | 0.831 | 0.813 | 0.849 | 0.066 | 0.652 | 0.885 | 0.777 |
| GSVA | 0.811 | 0.793 | 0.830 | -0.137 | 0.729 | 0.751 | 0.741 |
| ***CellCycle*** | | | | | | | |
| SuperFeat | 0.996 | 0.994 | 0.998 | 0.062 | 0.964 | 1.000 | 0.965 |
| Seurat | 0.954 | 0.921 | 0.987 | 0.057 | 0.977 | 0.818 | 0.973 |
| singscore | 0.979 | 0.964 | 0.993 | 0.038 | 0.888 | 0.964 | 0.891 |
| AUCell | 0.930 | 0.890 | 0.969 | 0.029 | 0.983 | 0.727 | 0.976 |
| GSVA | 0.942 | 0.902 | 0.982 | 0.148 | 0.978 | 0.782 | 0.973 |

**Table S2.** The datasets for feature training

| State | Type | Dataset | AccessionID | PMID | nCell | CellType |
|---|---|---|---|---|---|---|
| Exhasution | train | KIRC | GSE111360 | 30550791 | 2,777 | CD8 T cells |
| Exhasution | validation | HCC | GSE98638 | 28622514 | 3,636 | T cells |
| EMT | train | PDAC | GSE154778 | 32988401 | 4,604 | Epithelial cells |
| EMT | validation | HGSOC | GSE132149 | 32049047 | 1,410 | Epithelial cells |
| Hypoxia | train | Glioma | GSE84465 | 29091775 | 915 | Neoplastic |
| Hypoxia | validation | Glioma | GSE131928 | 31327527 | 7,062 | Malignant |
| CellCycle | train | LIHC | GSE140228 | 31675496 | 3,020 | T cells |
| CellCycle | validation | UCEC | GSE139555 | 32103181 | 12,692 | T cells |
| CellCycle | validation | BRCA | GSE110686 | 29942092 | 5,987 | T cells |
| Quiescent | train | CRC | GSE163974 | 33898197 | 218 | Cancer Stem Cells |
| Differentiation | train | LUNG | GSE135893 | 32832598 | 16,033 | Ciliated cells |
| Angiogenesis | train | CRC | GSE146771 | 32302573 | 2,155 | Macrophage |
| MacM1Polarization | train | KIRC | GSE111360 | 30550791 | 9,746 | Macrophage |

| | | | | | | |
|---|---|---|---|---|---|---|
| MacM2Polarization | train | KIRC | GSE111360 | 30550791 | 9,746 | Macrophage |
| Inflammation | train | CDC | GSE154763 | 33545035 | 5,461 | Dendritic cells |
| vCAFSignature | train | ICC | GSE142784 | 32505533 | 6,475 | Fibroblast |
| mCAFSignature | train | ICC | GSE142784 | 32505533 | 6,475 | Fibroblast |
| apCAFSignature | train | ICC | GSE142784 | 32505533 | 6,475 | Fibroblast |
| iCAFSignature | train | ICC | GSE142784 | 32505533 | 6,475 | Fibroblast |
| progenitorCAF | train | Melanoma | submission | submission | 4,594 | Fibroblast |
| progenitorCAF | test | FibroXplorer | Multiple-source | 33981032 | 99,596 | Fibroblast |

21
